# Deep origin of the long root tuft: the oldest stalk-bearing sponge from the Cambrian Stage 3 black shale of South China

**DOI:** 10.1101/2023.04.07.535994

**Authors:** Hao Yun, Dorte Janussen, Xingliang Zhang, Joachim Reitner

## Abstract

In virtue of remarkable tolerance on hypoxia and adaptive specialization in morphology, diverse hexactinellid sponges were prosperous in an early Cambrian living condition that characterized by dysoxic to anoxic bottom waters documented by black shales. New fossils from the black shale of Niutitang Formation (basal Stage 3 of Cambrian) in Hunan Province of China, reveal for the first time an articulated body of the sponge *Hyalosinica archaica* Mehl & Reitner in Steiner *et al*., 1993, which possesses an ovoid main body and an impressive long stalk. The specular skeleton includes large diactines that are generally organized as fan-shaped clusters, a few small stauractines and hexactines, and twisted bundles of long monaxons that form the stalk/root tuft. This hexactinellid sponge represents the oldest extinct taxon that took advantage of a long stalk to elevate the main body above the sediment surface and thus to adapt to the oxygen-deficient sea-bottom environment. The long root tuft links *Hyalosinica* to a series of fossil and recent sponge taxa and proves a deep origin of the stalk-bearing morphology, indicating a likely parallel evolution within the Hexactinellida in response to special environmental pressures. Furthermore, the overall skeletal organization indicates that *Hyalosinica*, as well as related early “rossellimorphs”, are basal stem group representatives of Hexactinellida and probably branched before the extinct Reticulosa and before the two extant hexactinellid subclasses.

## Introduction

Sponges are a group of basal animals significant for deciphering the origin and early evolution of metazoans (e.g. Ax 1996; Reitner & Mehl 1996; Mehl-Janussen 1999; Philippe *et al*. 2009; Dohrmann & Wörheide 2013). However, related paleontological investigations are inhibited by the incomplete fossil record from the early time. Since most of the Precambrian sponge-like fossils are more or less questionable (Gehling & Rigby 1996; Brasier *et al*. 1997; Li *et al*. 1998; Reitner & Wörheide 2002; Clapham *et al*. 2004; Clites *et al*. 2012; Antcliffe *et al*. 2014; Turner 2021), the earliest sponges (spicule fossils) have been dated back to the beginning of Cambrian, *ca*. 535 Ma (Chang *et al*. 2017, 2019; Reitner *et al*. 2022). Probable stem-group hexactinellids and demosponges were revealed by skeletal frames from the Cambrian Stage 2 phosphorites of the basal Niutitang Formation in Hunan, China (Luo & Reitner 2019) and the upper Pusa Formation in Spain (Reitner *et al*. 2022). The earliest diverse articulated body fossils of sponges, which could shed more light on the evolution of sponge morphology and adaptive strategies, were discovered from the Cambrian Stage 3 Burgess Shale-type Lagerstätten (e.g. Rigby & Hou 1995; Wu *et al*. 2014; Botting & Peel 2016; Luo *et al*. 2020), and noticeably, fossil assemblages have been found in the early Cambrian black shales (Steiner *et al*. 1993; Yuan *et al*. 2002; Xiao *et al*. 2005; Yang *et al*. 2010; Yun *et al*. 2022).

Even though the Cambrian black shales mostly represent a shelf margin to slope environment with dysoxic to anoxic bottom waters that is supposed to be inhospitable for most benthic animals (Zhou & Jiang 2009; Jin *et al*. 2016; Yeasmin *et al*. 2017; Zhang 2022), sponge communities (with a relatively high diversity of hexactinellids) were prosperous in these conditions due to their remarkable tolerance on hypoxia (Leys & Kahn 2018), adaptive specialization of morphology and life style, and mutualistic interactions with anaerobic microbes (Tabachnick 1991; Hoffmann *et al*. 2005; Mills *et al*. 2014; Tatzel *et al*. 2017; Morganti *et al*. 2021; Micaroni *et al*. 2022; Yun *et al*. 2022). More than 17 genera and 21 species of sponges were restored from the skeletal frameworks that were discovered in the black shales of Niutitang, Shuijingtuo, Hetang, and Huangboling formations in the South China block (Steiner *et al*. 1993; Mehl & Erdtmann 1994; Wu *et al*. 2005; Xiao *et al*. 2005; Yang *et al*. 2010; Luo *et al*. 2021). Within these taxa, *Hyalosinica archaica* Mehl & Reitner in Steiner *et al*., 1993 was regarded as the oldest known root tuft or stalk of a hexactinellid sponge (Steiner *et al*. 1993). However, there was no evidence for the main body of the stalk due to the absence of complete body fossils (Steiner *et al*. 1993; Xiao *et al*. 2005). Furthermore, *Solactiniella plumata* Mehl & Reitner in Steiner *et al*., 1993 was described based on fossils with unclear arrangement of spicules and incomplete bodies, and was usually assigned to a class-uncertain group (Steiner *et al*. 1993; Xiao *et al*. 2005; Yang *et al*. 2010). Therefore, it is necessary to explore new fossil materials to reconstruct the overall body of the root tuft-bearing sponges and to decipher the taxonomy and evolution of these enigmatic early sponge taxa.

New fossils from the Niutitang Formation (basal Stage 3 of Cambrian) in the Sancha section of Hunan Province, China, reveal complete bodies of the long stalk-bearing sponge and suggest that *H. archaica* and *S. plumata* were came from a same hexactinellid species. The distinctive long root tuft, as well as the spicule arrangement pattern, of this sponge are significant in interpreting the adaptive strategy and phylogenetic relationship of early hexactinellids.

## Material and methods

The specimens were collected from the lower Niutitang Formation (basal Stage 3 of Cambrian) in the Sancha section of Zhangjiajie, Hunan Province, China. The geological setting and related geochemical analysis are well described in Steiner *et al*. (1993) and Steiner *et al*. (2001). The holotypes of *Hyalosinica archaica* (San109A, B) and *Solactiniella plumata* (San101 A, B) from the Sancha section were originally described by Mehl & Reitner in Steiner *et al*. (1993). New materials studied herein include two specimens preserved with relatively complete bodies of the stalk-bearing sponge and four with fragments (see Supplementary material for the specimen list). All specimens are deposited as palaeontology collections at the Geobiology Department and Geoscience Museum of the University of Göttingen.

Fossils were examined and prepared under a Zeiss Stemi2000 stereomicroscope, and photographed with a Sony Alpha 7Ⅳ camera under the magnesium light in the Geosciences Center of the University of Göttingen. The images were re-levelled and processed with Affinity Photo 2.0 and CorelDRAW X8. X-Ray Fluorescence (XRF) analysis was performed under a Bruker’s M4 TORNADO spectrometer (Geosciences Center of the University of Göttingen) to investigate the elemental composition of the fossils.

## Results

### Fossil preservation

The fossils are mainly preserved as solid internal moulds and/or hollow impressions of spicules (Figs 1–3). The XRF analysis shows that there are strong signals of the elements Fe, K, Si, Al, and Ti; specifically, Fe is more concentrated in the spicules (Fig. 1C, D; Supplementary material). This result indicates a composition of iron oxides and clayey minerals, and the fossils were experienced a secondary silicification and pyritization. Furthermore, the fossils are goldish and grey in colour due to the long-time weathering and oxidation after exposure to the air. Although the soft part is not preserved, the pyritized spicules can well reflect the original skeletal architecture and framework pattern of the sponge.

**Fig. 1.**
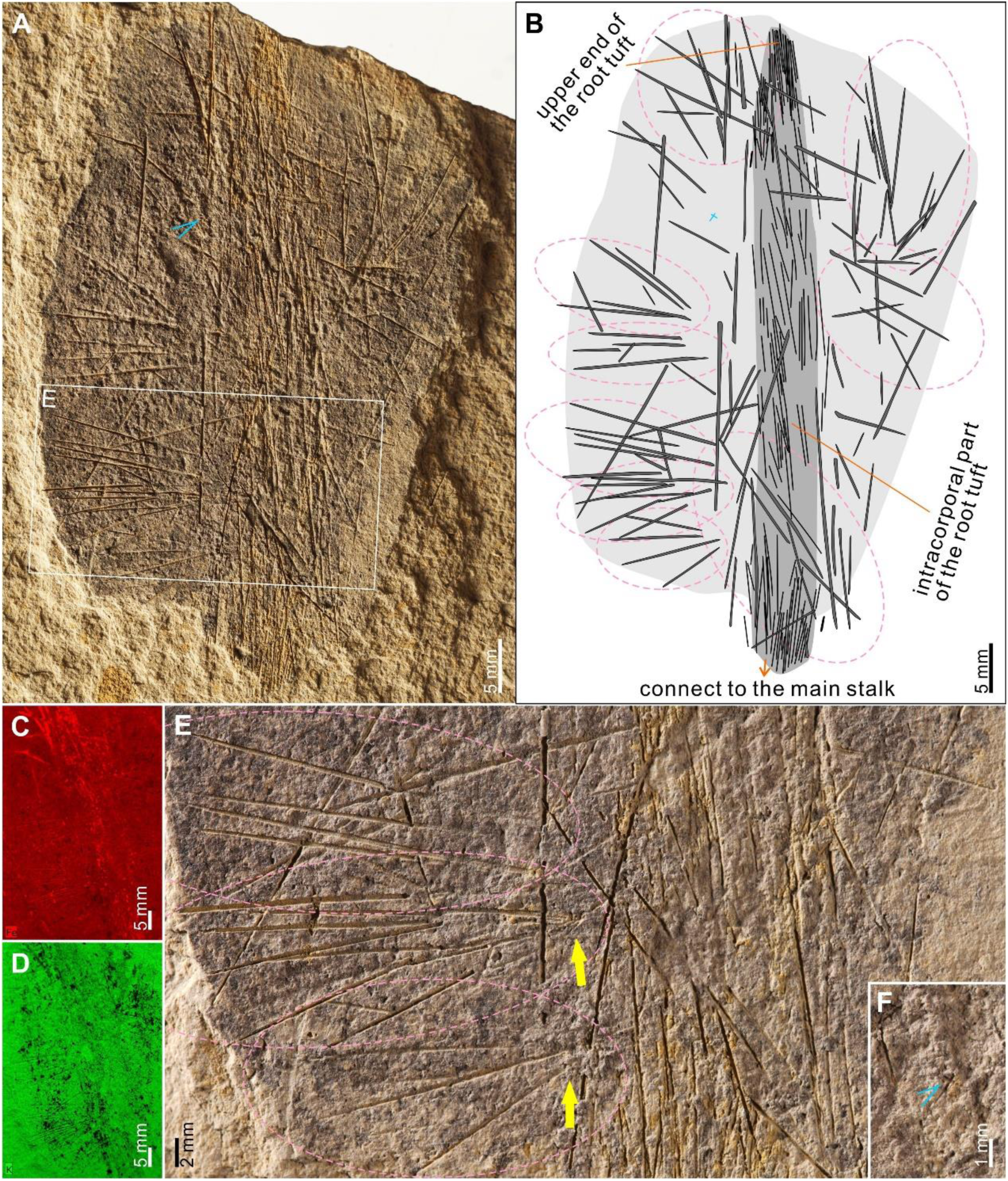
Body structure of *Hyalosinica archaica* from the Niutitang Formation (Cambrian Stage 3) in the Sancha section, Hunan Province, China. A: ovoid body and intracorporal part of the stalk distributed along the central axis of the body, specimen (GZG INV. 998). B: interpretation of the skeletal structure. C, D: representative XRF images of the specimen (GZG INV. 998), showing the concentration of elements Fe (upper) and K (lower). E: detailed view of the box indicated in (A), showing fan-shaped diactine clusters (indicated by pink dotted-line circles, and their converging ends indicated by yellow arrows). D: a small diactine (indicated by a light blue marker) in the main body.

### The complete body of *Hyalosinica archaica*

New fossil specimens reveal that the overall body of *H. archaica* consists of two parts: an ovoid upper part (the main body) and a long root tuft that originally inserted upward into the main body and anchored downward to the seafloor with a relatively loose end (basal end).

The specimen MSan385 represents an ovoid sponge body possessing an upper part (intracorporal part) of the root tuft that is situated at the axial area of the body (Fig. 1A). The main body is 33.7 mm in maximum width and 45.5 mm in height, while the tuft at the central axis is 54 mm high, 2.6 mm wide at its upper end, and 4.3 mm wide at the lower end. The lower end protrudes out the body to form a long stalk. The specimen MSan383 shows a long extracorporeal root tuft (stalk) along with a lower part of the ovoid body, in which the root tuft is 41.4 mm high, and 4.1 mm (close to the basal end) tapering to 2.5 mm (proximal end) in width. The original height of the main body of this specimen is estimated to be more than 22 mm based on the partly preserved body part (Fig. 2A).

**Fig. 2.**
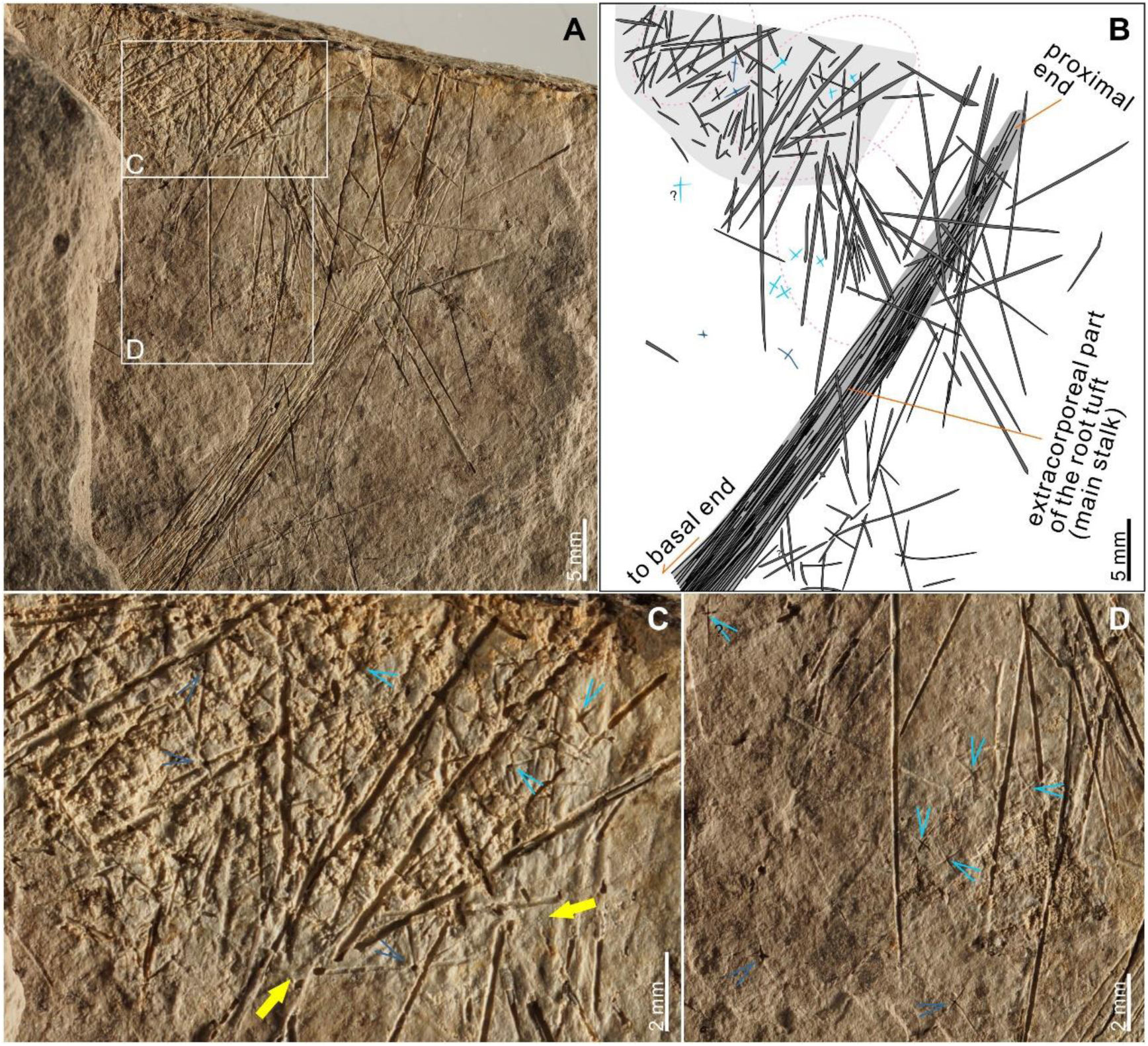
Long stalk and lower part body of *Hyalosinica archaica* from the Niutitang Formation (Cambrian Stage 3) in the Sancha section, Hunan Province, China. A: part of the ovoid body along with its extracorporeal long stalk, specimen (GZG INV. 999) B: Interpretation of the skeletal structure and the stalk; fan-shaped diactine clusters are indicated by pink dotted-line circles. C, D: detailed view of boxes in (A), showing fan-shaped diactine clusters (the converging ends are indicated by yellow arrows) and small diactines (indicated by light blue markers) and hexactines (and probably pentactines; indicated by dark blue markers) within and exfoliated from the main body.

### Spicule structure and arrangement

Spicules in the main body of *H. archaica* are dominated by large diactines (and possibly monactines) that range from 7 mm to 21 mm in length and 0.08 mm to 0.30 mm in diameter (Figs 1–3). The diactines are organized as fan-shaped clusters and distributed around the central axial tuft (Figs 1B, 3B). In each cluster, there are three to five spicules with one end converging together and the other end gradually diverging outwards (Fig. 1B, E). The converging ends of the clusters are distributed along the longitudinal central axis while the diverging ends radiate towards and project from the margin of the sponge body (Fig. 1B). However, in many specimens, the diactines are interlaced to each other and the organization seem irregular due to the compression during burial and incomplete preservation (Fig. 3A, B). Furthermore, in some specimens, there are a few small diactines, stauractines, and hexactines (and possible pentactines), with length less than 1.5 mm and diameter less than 0.06 mm, scattered in the interspace of the large diactines (Figs 1A, F, 2, B, C).

**Fig. 3.**
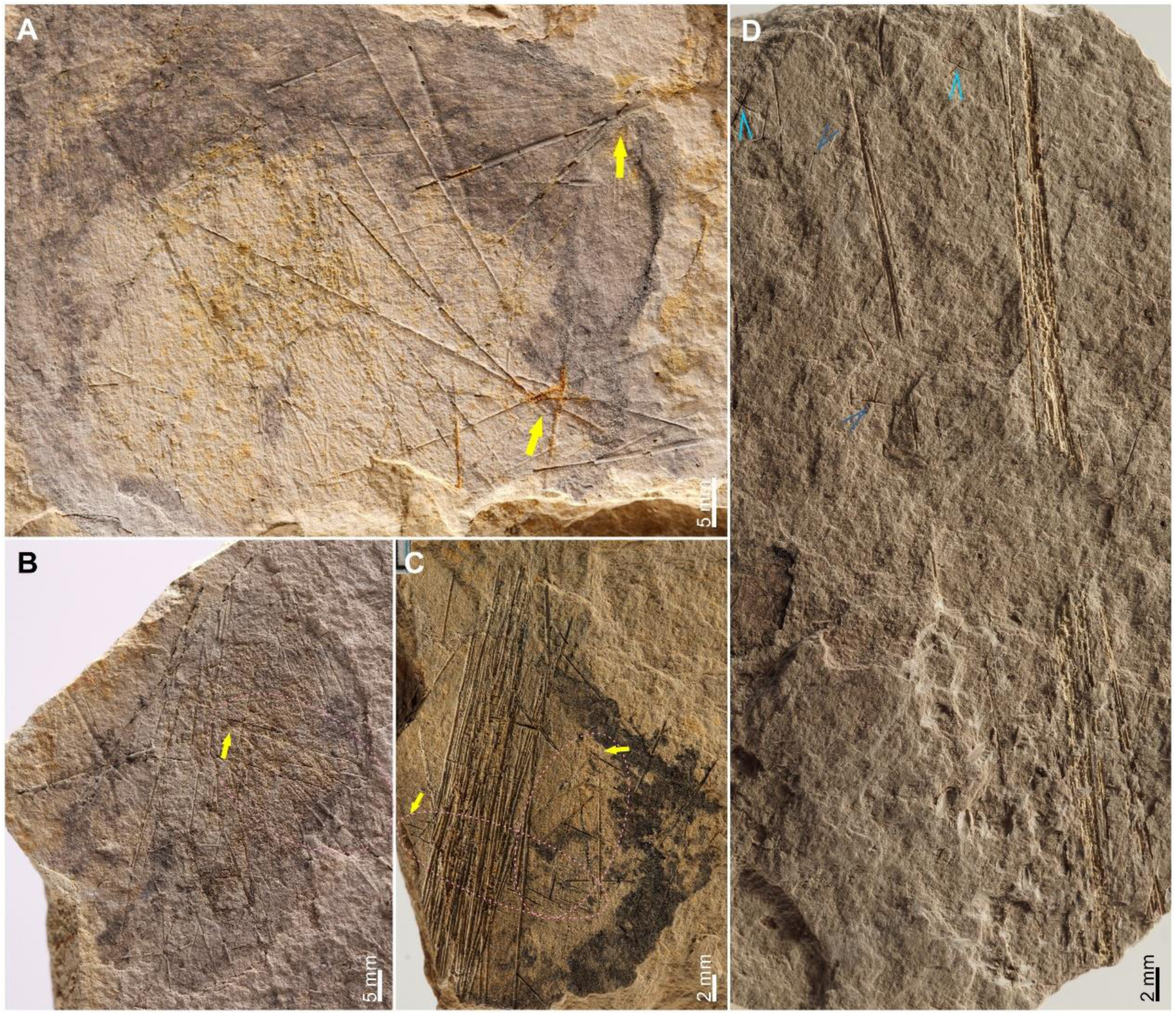
Other representative specimens of *Hyalosinica archaica* from the Niutitang Formation (Cambrian Stage 3) in the Sancha section, Hunan Province, China. A, B: fragments of ovoid bodies, specimens (GZG INV. 1000) (A) and (GZG 1001) (B). C, D: fragments of the long root tuft (stalk), specimens (GZG INV. 1002) (C) and (GZG INV. 1003) (D). The fan-shaped diactine clusters are indicated by pink dotted-line circles while their converging ends are indicated by yellow arrows; small diactines and hexactines (and probably pentactines) exfoliated from the main body and surrounding the proximal part of the stalk are indicated by bright blue and dark blue markers, respectively.

Spicules of the root tuft are bundled, long diactines and probably monactines (Figs 2A, 3C, D). The length of the diactines reaches 22 mm and the average diameter is 0.1 mm in paratype MSan383 (Fig. 2A), though they can be thicker (0.25 mm in diameter) in some fragmental specimens such as MSan384 (Fig. 3C). The spicules are compact and parallel to each other in the bundles, while the bundles are twistedly organized and interwoven (roughly in a clockwise direction if observed from the top) to form the tuft (Figs 1B, 2B). There are coarse diactines, some organized within fan-shaped clusters, and small diactines and triaxons perpendicular to or crossing with the root tuft, especially in the fragmental specimens (Figs 2D, 3C, D), which likely represent parenchymal spicules exfoliated from the ovoid main body instead of *in-situ* structures.

## Discussion

### Skeletal architecture

The skeleton of *Hyalosinica* consists of three main groups of spicules: parenchymal principalia, parenchymal intermedia, and basalia (Fig. 4). The principalia are large diactines organized in clusters and roughly radiate from the vertical central axis to the outer surface (and the tips could protrude out the surface) of the ovoid body. The intermedia, including a few small diactines, stauractines, and hexactines, are scattered in the interspace of the framework formed by the principalia. The basalia are characterized by long diactines and maybe monactines that bundled together and nearly parallel to each other; the bundles are twisted in a clockwise orientation and intertwined to form a long, sturdy root tuft.

**Fig. 4.**
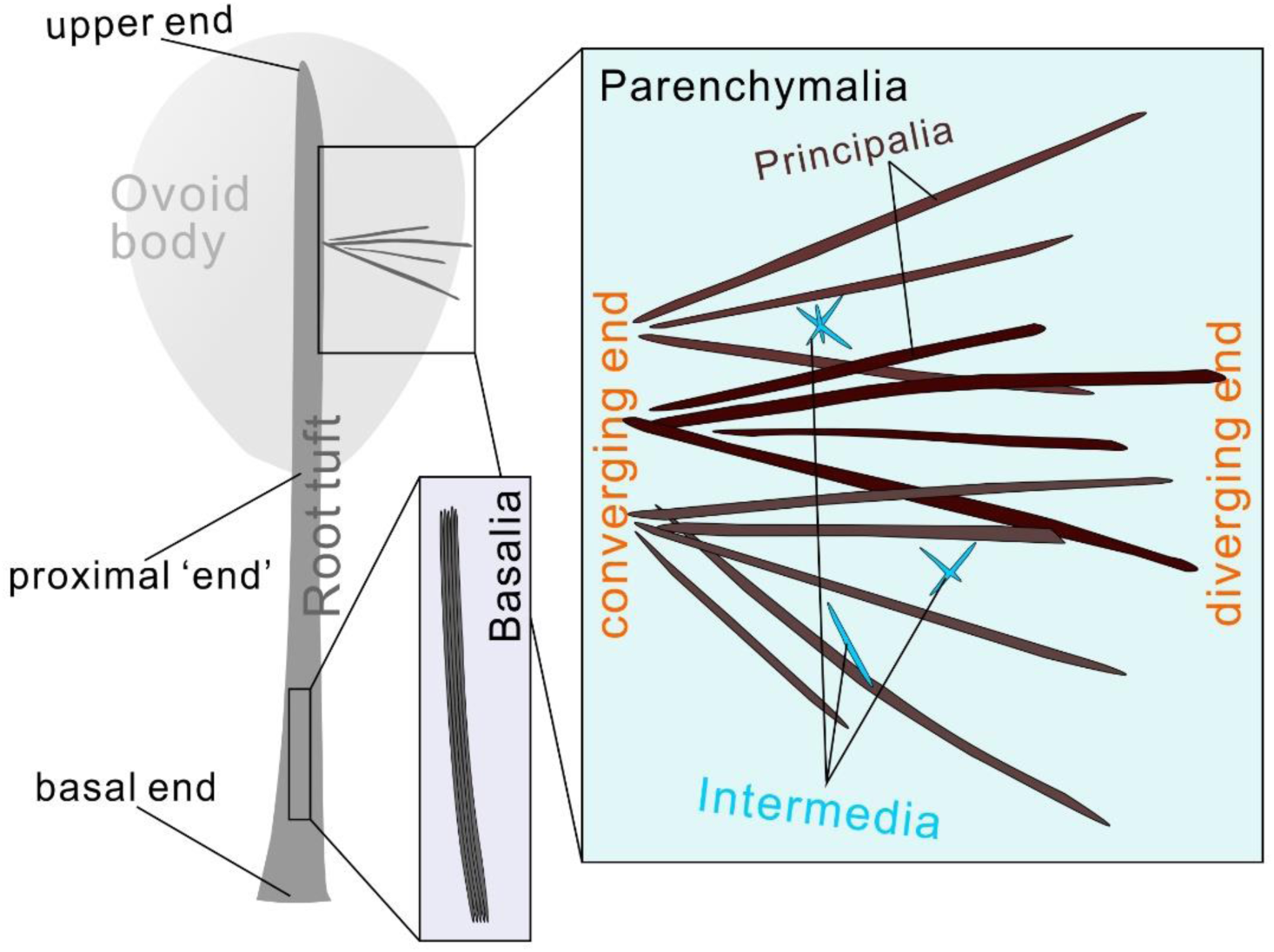
Overall structure and spicule arrangement of *Hyalosinica archaica* from the Niutitang Formation (Cambrian Stage 3) in the Sancha section, Hunan Province, China. The body is composed of an ovoid body and a long root tuft (stalk). The spicules include parenchymal principalia, parenchymal intermedia, and basalia; the large diactines (principalia) are generally organized as fan-shaped clusters.

The structure of large diactine clusters is characteristic in *Hyalosinica*. These clusters are fan-shaped in compressed fossils (thus possibly in subconical shape originally), each composed of mostly three to five diactine spicules. Spicule tips in the converging end of each cluster are tightly contacted (even partly fused together) (Figs 1C, 3B), or just loosely intersected (Figs 2C, 3A, C). The clusters, especially those with tightly contacted spicules, likely functioned in supporting and strengthening the main body, reminiscent of the large dermal pentactines in the coeval sponge *Sanshapentella* (Mehl & Erdtmann 1994; Luo *et al*. 2021; Yun *et al*. 2022). However, the diactine clusters herein are distinct from the dermal pentactines both in detailed structure and orientation, since the large dermal pentactines of *Sanshapentella* are composed of four long rays inserting into the body and a tiny fifth ray protruding out the body (Yun *et al*. 2022).

The root tuft of *Hyalosinica* can be subdivided into two parts: a long extracorporeal stalk that originally anchored to the seafloor with a loose basal end and an intracorporal part that deeply inserted into the main body (Figs 4, 5), similarly to the modern genus *Hyalonema* Gray, 1832 (Hexactinellida: Amphidiscophora: Hyalonematidae). The intracorporal part of the root tuft was likely beneficial in reinforcing the connection between the stalk and the main body to prevent breaking, since the long stalk is much thinner than the main body (the average ratio of diameters of the stalk and the ovoid body is about one eighth).

**Fig. 5.**
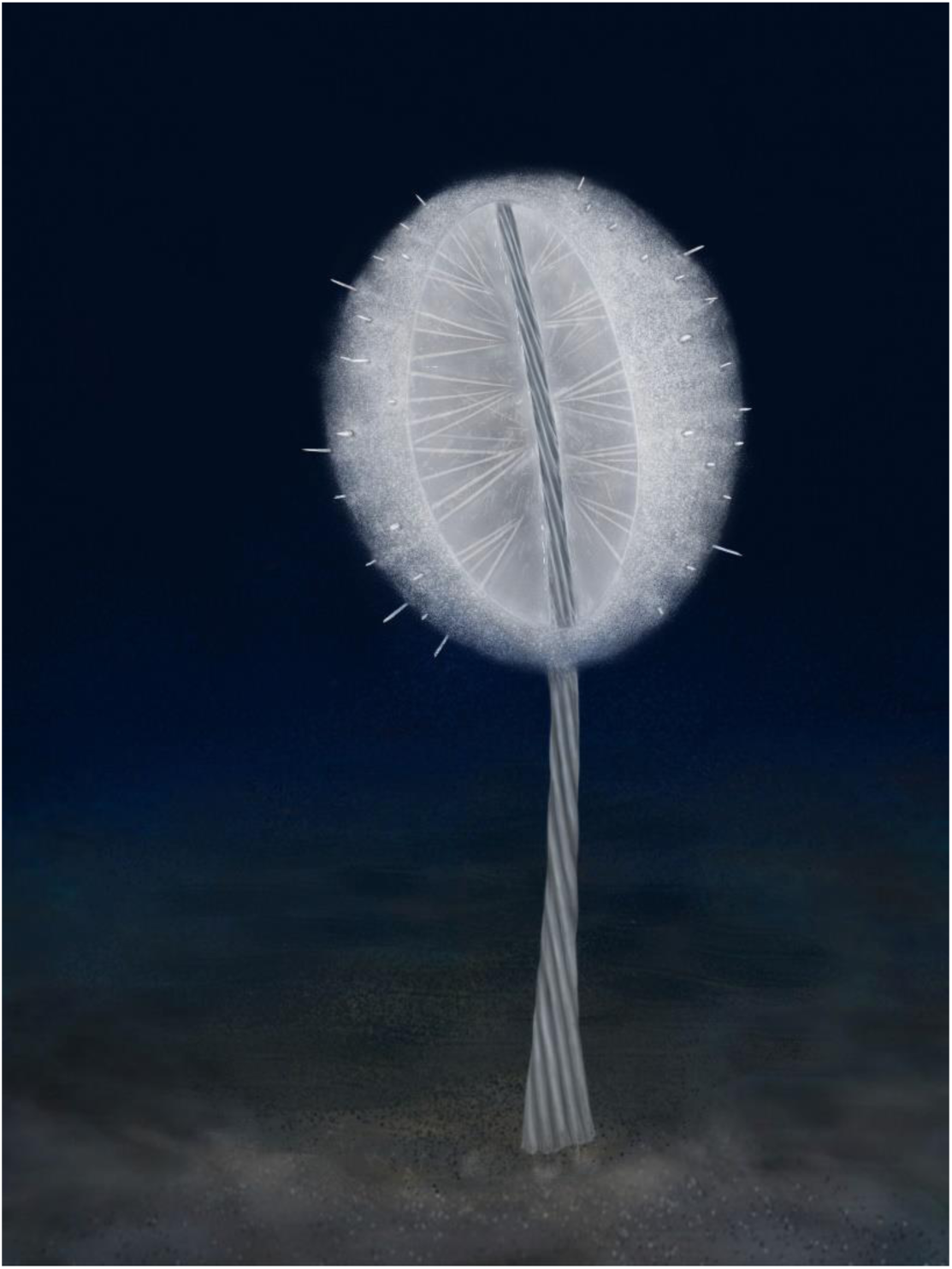
Artistic reconstruction of *Hyalosinica archaica* from the Niutitang Formation (Cambrian Stage 3) in the Sancha section, Hunan Province, China.

### Long root tuft (stalk): an adaptive strategy

The long root tuft of *Hyalosinica* is reminiscent of the modern hexactinellid *Hyalonema*, which has a flexible stalk (can be more than one meter long) formed by spirally twisted monactines and associated diactines (Schulze 1887; Tabachnick & Menshenina 2002). There are also a series of fossil hexactinellid sponges possessing conspicuous root tufts and/or long stalks, including some species of *Protospongia* Salter, 1864 from the middle Cambrian of Quebec, Canada (Dawson & Hinde 1888, 1889), *Retifungus rudens* Rietschel, 1970 from the Lower Devonian Hunsrück Slate of Germany (Rietschel 1970), *Hyalostelia smithii* (Young & Young, 1877) from the Mississippian (lower Carboniferous) of Ayrshire, Scotland (Young & Young 1877; Reid 1968), *Stioderma coscinum* Finks, 1960 from the Pennsylvanian (upper Carboniferous) to Permian of southwestern United States (Finks 1960), *Hyalonema cretacea* Mehl & Hauschke, 1995 from the Campanian Stage, Upper Cretaceous of northwest Germany (Mehl & Hauschke 1995), and *Hyalonema vetteri* Janussen, 2014 from the Coniacian Stage, Upper Cretaceous of the isle Bornholm, Denmark (Janussen 2014). Each of these root tufts is subtly different from all the others (Table 1): the recent and fossil *Hyalonema*, as well as *Hyalosinica* described herein, have slender but sturdy stalks formed by long, twisted bundles of diactines and possibly monactines; the three late Palaeozoic taxa, *Stioderma*, *Hyalostelia* and *Retifungus*, have more flexible tufts with monaxon spicules slightly twisted or not twisted at all; the tuft of *Stioderma* is formed by a bundle of parallel, smooth monaxons and thus can be relatively thick in diameter; while the “stalk” of *Protospongia* is just a few of long basal spicules instead of a real tuft, faintly reminiscent of the modern genus *Monorhaphis* Schulze, 1904 (Hexactinellida: Amphidiscophora: Monorhaphididae). Nevertheless, all these root tufts or stalks, are as long as, or much longer than their main bodies and were positioned between the seafloor and the main body at the top. Therefore, similarly to the life mode of recent *Hyalonema*, all of them functioned by anchoring the sponge firmly to the sea floor and elevating the suspension feeding main part of the body well above the substrate (Gray 1869; Schulze 1886, 1887; Beaulieu 2001; Tabachnick & Menshenina 2002).

**Table 1.**
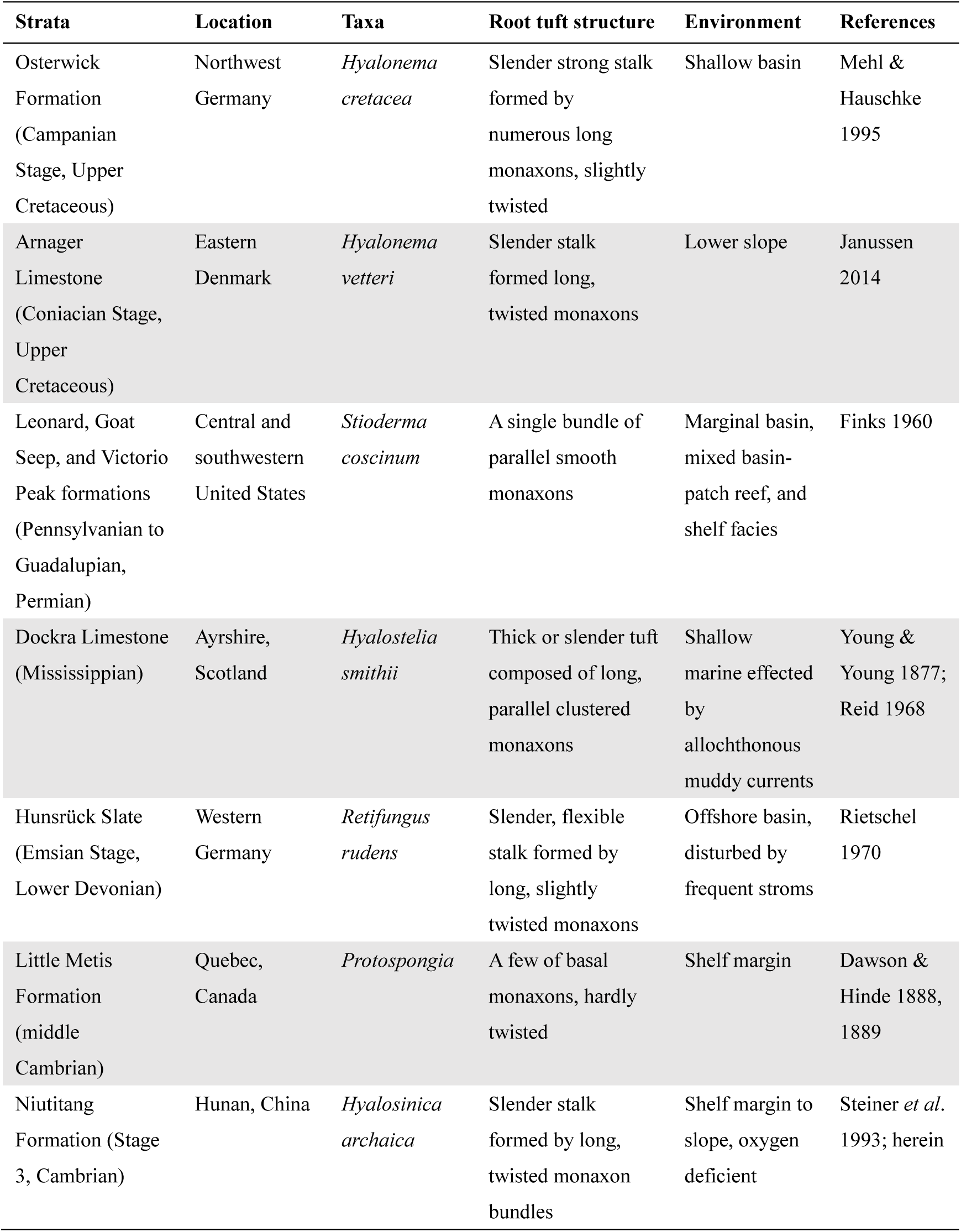
Representative fossil hexactinellids with a long root tuft.

| Strata | Location | Taxa | Root tuft structure | Environment | References |
| --- | --- | --- | --- | --- | --- |
| Osterwick Formation (Campanian Stage, Upper Cretaceous) | Northwest Germany | *Hyalonema cretacea* | Slender strong stalk formed by numerous long monaxons, slightly twisted | Shallow basin | Mehl & Hauschke 1995 |
| Arnager Limestone (Coniacian Stage, Upper Cretaceous) | Eastern Denmark | *Hyalonema vetteri* | Slender stalk formed long, twisted monaxons | Lower slope | Janussen 2014 |
| Leonard, Goat Seep, and Victorio Peak formations (Pennsylvanian to Guadalupian, Permian) | Central and southwestern United States | *Stioderma coscinum* | A single bundle of parallel smooth monaxons | Marginal basin, mixed basin-patch reef, and shelf facies | Finks 1960 |
| Dockra Limestone (Mississippian) | Ayrshire, Scotland | *Hyalostelia smithii* | Thick or slender tuft composed of long, parallel clustered monaxons | Shallow marine effected by allochthonous muddy currents | Young & Young 1877; Reid 1968 |
| Hunsrück Slate (Emsian Stage, Lower Devonian) | Western Germany | *Retifungus rudens* | Slender, flexible stalk formed by long, slightly twisted monaxons | Offshore basin, disturbed by frequent stroms | Rietschel 1970 |
| Little Metis Formation (middle Cambrian) | Quebec, Canada | *Protospongia* | A few of basal monaxons, hardly twisted | Shelf margin | Dawson & Hinde 1888, 1889 |
| Niutitang Formation (Stage 3, Cambrian) | Hunan, China | *Hyalosinica archaica* | Slender stalk formed by long, twisted monaxon bundles | Shelf margin to slope, oxygen deficient | Steiner *et al.* 1993; herein |

In general, the long root tuft help sponges adapting to two main types of environmental parameters. The first type is a habitat exposed to frequent currents from different directions. With a flexible root tuft, sponges such as *Hyalostelia* and *Retifungus* can always regulate their osculum and canal system to fit with the water current, thereby aiding their filtering nutrition from the surrounding seawater (Mehl 1996). The second type is a condition with anoxic or dysoxic, commonly soft substrates, dominant especially in the deep sea. The root tuft of sponges, such as *Hyalonema* and *Hyalosinica* described herein, can anchor the sponge to soft sediments in lack of hard substrate, and a long stalk can keep the body at a certain height above the sediment surface, which is particularly advantageous in the case of dysoxia and/or fine-grained sedimentary input (Janussen 2014).

A series of geochemical analyses on redox-sensitive trace elements, highly reactive iron speciation, and pyrite S-isotopic compositions revealed a stratified oceanic paleoenvironment during the sponge-bearing interval of the Cambrian Niutitang Formation, in which oxic surficial water was underlain by anoxic or euxinic deep water (e.g. Wang *et al*. 2015; Jin *et al*. 2016; Yeasmin *et al*. 2017). These studies also suggested that a small amount of oxygen might be episodically introduced to the ferruginous- or euxinic-dominated bottom water by currents and/or hydrothermal events, forming a localized condition with oxygen level above the lethal threshold for sponges but still inhospitable to most epifauna or nektons (Steiner *et al*. 2001; Zhou & Jiang 2009; Cheng *et al*. 2017). Besides the paleoenvironmental view, adaptive strategies exploited by sponges themselves, such as a morphology with noticeable projections (functioned in enlarging the relative surface area; Xiao *et al*. 2005; Yun *et al*. 2022) and a long stalk (this study), are also a significant reason for the development of the black shale sponge community. Specifically, it is reasonable to assume that the long stalk enabled *Hyalosinica* to reach the oxic (at least not euxinic) water. Therefore, of all the stalk-bearing sponges *Hyalosinica* represents the oldest taxon that developed and took advantage of a long stalk to adapt to the deep-water, soft bottom, and oxygen-deficient environment.

### Phylogenetic implications

Specimens from the same section (Sancha) that were previously attributed to *Hyalosinica* (root tufts composed of parallel, or slightly twisted long monaxons) and *Solactiniella* (spherical body dominated by diactines) are here suggested to be incomplete body parts of a same taxon. Therefore, the genus *Solactiniella* is considered as a junior synonym of *Hyalosinica*, and the diagnosis of *Hyalosinica* is herein emended and expanded to describe the complete sponge body (see Supplementary material for the systematic palaeontology). However, since early Cambrian articulated sponge fossils are rare and commonly incompletely preserved, for practical reasons (and for the work of future sponge palaeontologists) we recommend to not abandon the name *Solactiniella*. It is expedient to retain *Solactiniella* as a subgeneric taxon to design incomplete specimens of *Hyalosinica* (fossils preserved with only the main body, but no root tuft). Thus, we can use the subgenus name *Hyalosinica* (*Solactiniella*) to describe a globular sponge body with irregular to plumose skeleton consisting mainly of diactines, with no trace of a root or stalk, nor any other basal attachment structure.

*Hyalosinica* (and former *Solactiniella*) from early Cambrian black shales was previously assigned to an undetermined class and tentatively to the Hexactinellida, since there was no definite evidence for clear triaxons in the skeletal framework (Steiner *et al*. 1993; Xiao *et al*. 2005; Yang *et al*. 2010). Stauractines and suspected hexactines were only discovered in *Solactiniella* cf. *plumata* from the Huangboling Formation (Cambrian Stage 4) of South China (Wu *et al*. 2005). The existence of certain parenchymal stauractines and hexactines in the new materials from the type locality finally concludes that this sponge was indeed a hexactinellid.

Although there is no consensus about the taxonomic or phylogenetic scheme of the early sponges (e.g., Kaesler 2004; Rigby & Collins 2004; Leys *et al*. 2007; Botting & Muir 2018), most articulated hexactinellids from Cambrian biotas are classified within the morphological group Reticulosa Reid, 1958 that characterized by a thin body wall and a series of parallelly-arranged stauractines, pentactines, and hexactines (Reid 1958; Rigby & Hou 1995; Finks & Rigby 2004; Rigby & Collins 2004; Yang *et al*. 2005; Wu *et al*. 2014). However, the body of *Hyalosinica* (originally described as *Solactiniella*) has a thick wall as well as inconspicuous oscula and a skeleton dominated by dense clusters of diactines that interlaced each other and organized in a generally plumose to radiate manner. Therefore, this genus is distinct from the typical Reticulosa and was instead classified into a supposedly polyphyletic group “Rossellimorpha” by Mehl (1996). The “Rossellimorpha” includes a number of fossil sponges with a thick body wall and an irregular distribution of reduced forms of hexactines (such as stauractines and diactines) (Mehl 1996). From this viewpoint, the skeleton of *Hyalosinica* represents an old, even basic, organization of the “Rossellimorpha”. Furthermore, the fan-shaped clusters of diactines and their general radiate arrangement show a degree of skeletal regulation, reflecting an organization within the Paleozoic taxa, which was dominated by highly irregularly-organized sponges (such as the nodular hexactinellid from the horizon just below the black shale of Niutitang Formation in the Sancha section, described by Luo & Reitner (2019)) as well as the Reticulosa with relatively well-organized skeletons.

As mentioned above, there are many fossil and recent hexactinellid sponges possessing conspicuous root tufts (Table 1). Despite the controversy regarding the taxonomy of the Devonian *Retifungus*, *Protospongia* (Cambrian to Devonian) is a typical Reticulosa, while *Hyalostelia* (Carboniferous) and *Stioderma* (Carboniferous to Permian) were classified into a superfamily Brachiospongioidea Finks, 1960 that was supposedly derived from the Reticulosa (Reid 1968; Mehl 1992, 1996; Krautter 2002; Finks & Rigby 2004). *Hyalonema* (late Cretaceous to present) belongs to the extant subclass Amphidiscophora Schulze, 1886, based on molecular data and types of their microscleres (amphidiscs and dermal triaxons) (Tabachnick & Menshenina 2002; Dohrmann *et al*. 2008). In absence of amphidiscs or related microscleres in the mentioned fossil sponges, it is still difficult to establish the phylogenetic relationship between Brachiospongioidea and Amphidiscophora. Hence, although the long root tuft links these fossil taxa to the recent *Hyalonema*, the stalk-bearing morphology more likely developed convergently (parallelly) within the Hexactinellida in response to special environmental pressures (Mehl 1996). A homoplasy within the subclass Hexasterophora Schulze, 1886 is the long stalk that characterizes the extant genus *Caulophacus* Schulze, 1886 (Hexactinellida: Hexasterophora: Rossellidae). According to molecular clock results, the subclasses Amphidiscophora and Hexasterophora diverged from a common ancestor during the middle to late Cambrian (Dohrmann *et al*. 2013). This ancestor may have derived from a “rossellimorph” taxon, which possessed key characters of both taxa, such as a long monaxonic stalk and a main body composed mainly of irregularly organized diactines and triaxons.

In summary, the body structure of *Hyalosinica* is characterized by a “rossellimorph” skeletal organization and a long stalk. Most likely parallelly, this type of stalk has evolved several times independently within the Hexactinellida, e.g. late Palaeozoic Brachiospongioidea and the recent *Hyalonema* (Fig. 6). Since the spicule arrangement of hexactinellids was likely evolved from irregular to regular (Mehl 1996; Reitner & Mehl 1996; Dohrmann *et al*. 2008; Luo & Reitner 2019), the reticulosan group branched off after the *Hyalosinica* and other basal “rossellimorphs” such as the Sancha nodular sponge (Luo & Reitner 2019) and *Sanshapentella* (Mehl & Erdtmann 1994; Yun *et al*. 2022). Furthermore, *Hyalosinica* represents the oldest known hexactinellids that developed an impressive stalk structure to colonize early Cambrian dysoxic marine environments, thus improving the adaptive potential and evolutionary success of the sponge phylum.

**Fig. 6.**
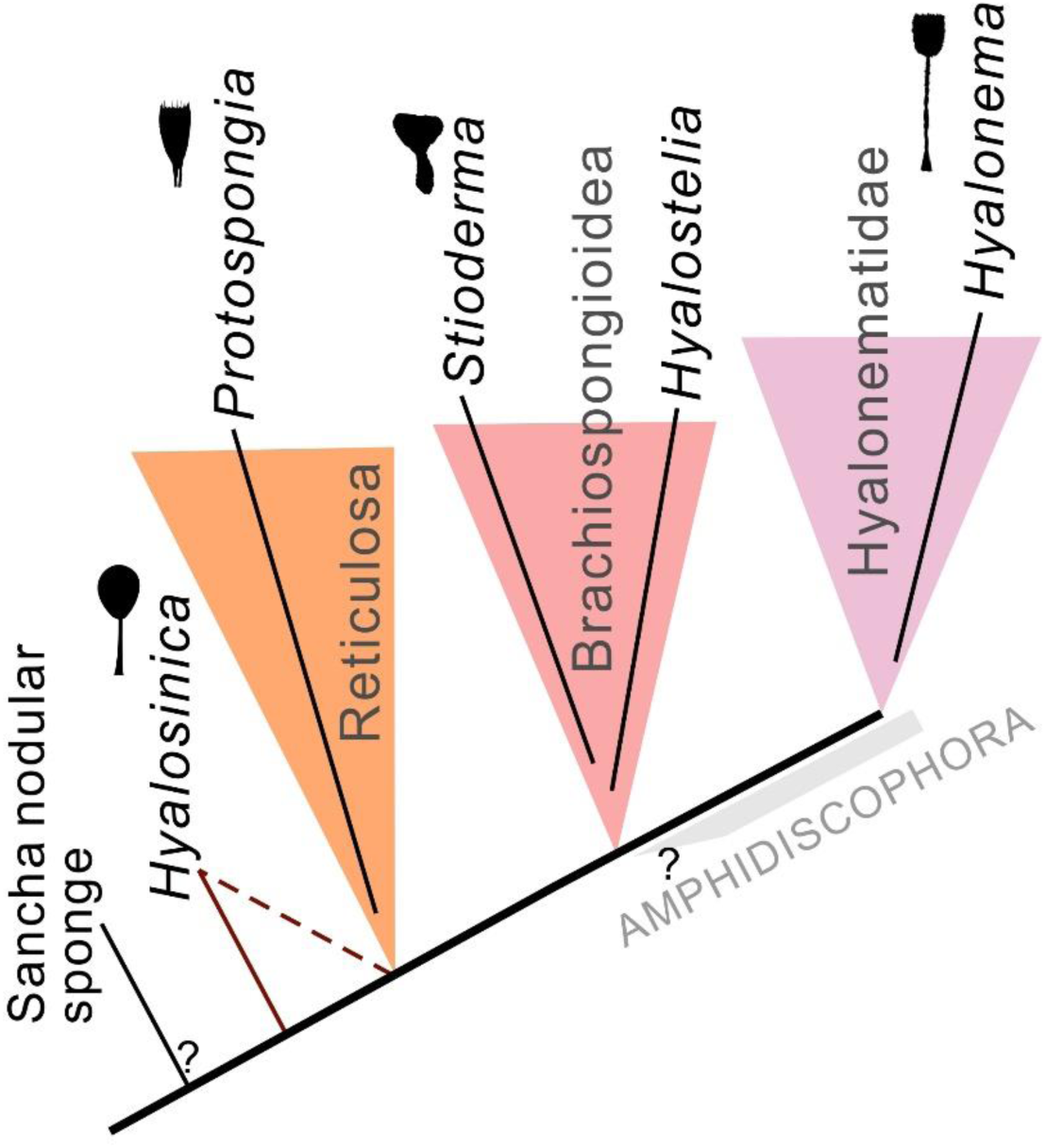
A possible phylogenetic scheme for the hexactinellid sponges with a long root tuft or stalk. The silhouettes of stalk-bearing sponges are drawn after the taxa listed in Table 1 (see Supplementary material for detail).

## Supporting information

1. Specimen list and original photographs 2. X-Ray Fluorescence (XRF) results of the specimen MSan385 3. Systematic palaeontology 4. Silhouettes of st

## Acknowledgments

We thank Gerhard Hundertmark (University of Göttingen, Centre of Geosciences) for assistance in specimen photographing, Xi Liu (Northwest University) for the artwork of fossil reconstruction, and Michael Steiner for providing some of the fossil specimens. This study was supported by the National Natural Science Foundation of China (grant nos. 42002011, 41890845, 41930319, and 41621003), Strategic Priority Research Program of Chinese Academy of Sciences (grant no. XDB26000000), and the 111 Project (grant nos. D17013). HY is sponsored by a Sino-German (CSC-DAAD) postdoc scholarship program (Ref. No. 202006970034/91809892). This study was supported by the Research Commission “Origin of Life” of the Göttingen Academy of Science and Humanities in Lower Saxony.

## Supplementary Material

The supplementary material can be found in the online version of the article. (*Here attached at the end of the manuscript)*

1. Specimen list and original photographs
2. X-Ray Fluorescence (XRF) results
3. Systematic palaeontology
4. Silhouettes of stalk-bearing sponges

