## Supplementary material for "Deep origin of the long root tuft: the oldest stalk-bearing sponge from the Cambrian Stage 3 black shale of South China": 1. Specimen list and original photographs 2. X-Ray Fluorescence (XRF) results of the specimen MSan385 3. Systematic palaeontology 4. Silhouettes of st

**This file includes:**

1. Specimen list and original photographs

2. X-Ray Fluorescence (XRF) results of the specimen MSan385

3. Systematic palaeontology

4. Silhouettes of stalk-bearing sponges

**1. Specimen list and original photographs**

*Specimen list*

GZG.INV.998 (main body + long root tuft)

GZG.INV.999 A, B (main body + long root tuft)

GZG.INV.1000 A, B (main body)

GZG.INV.1001 A, B (main body)

GZG.INV.1002 A, B (root tuft)

GZG.INV.1003 A, B (root tuft)

GZG.INV.999 A

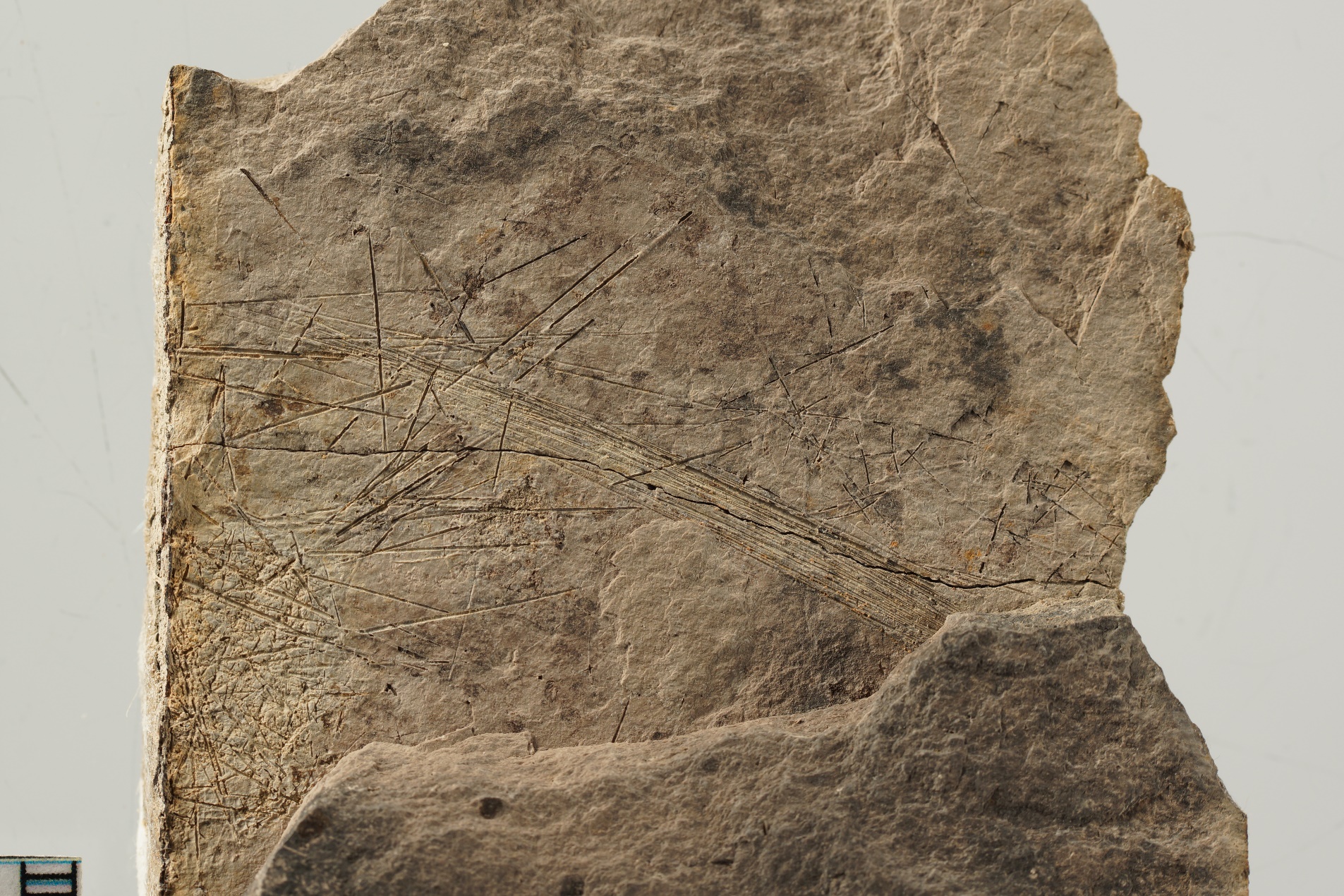

GZG.INV.999 B

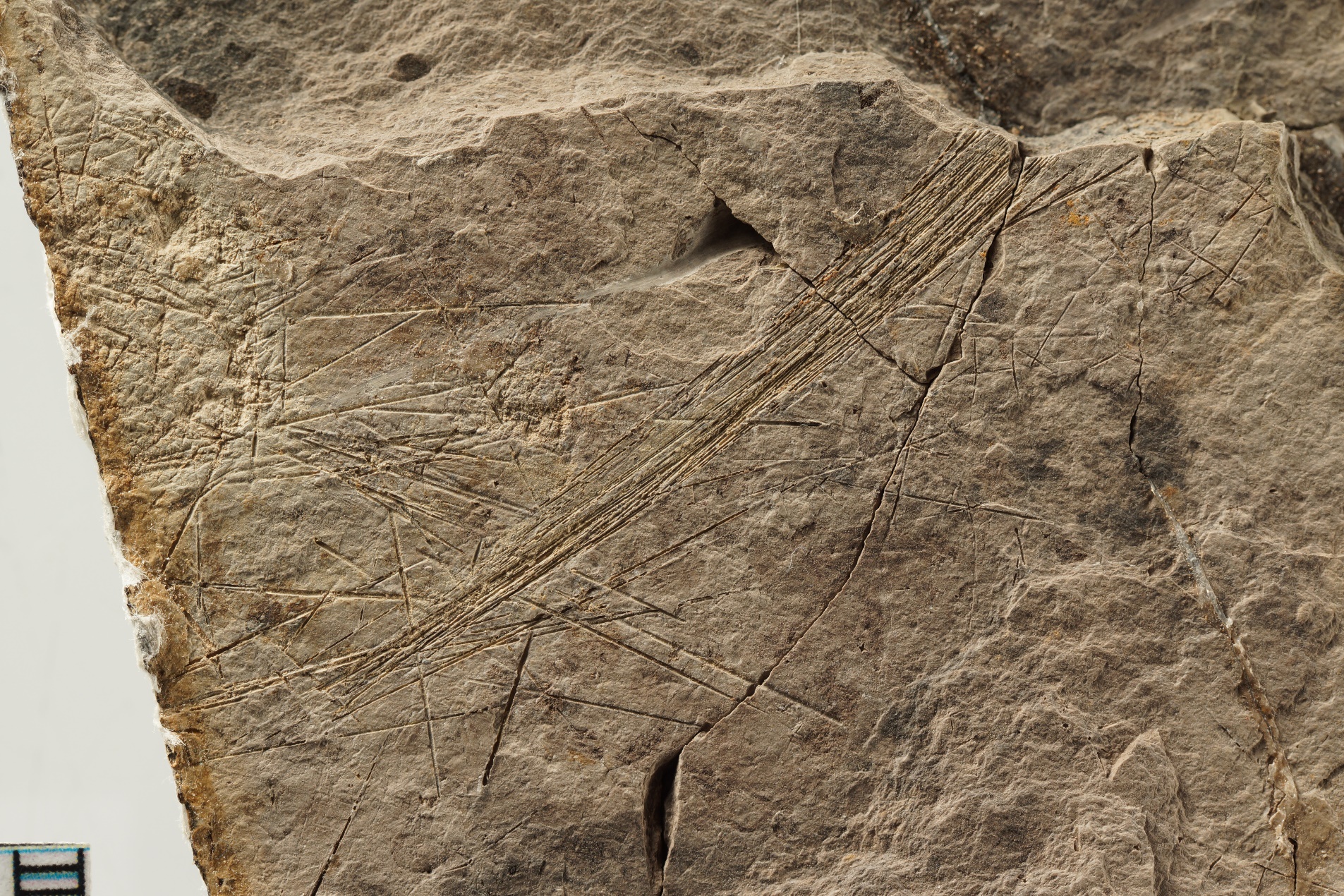

GZG.INV.1002 A

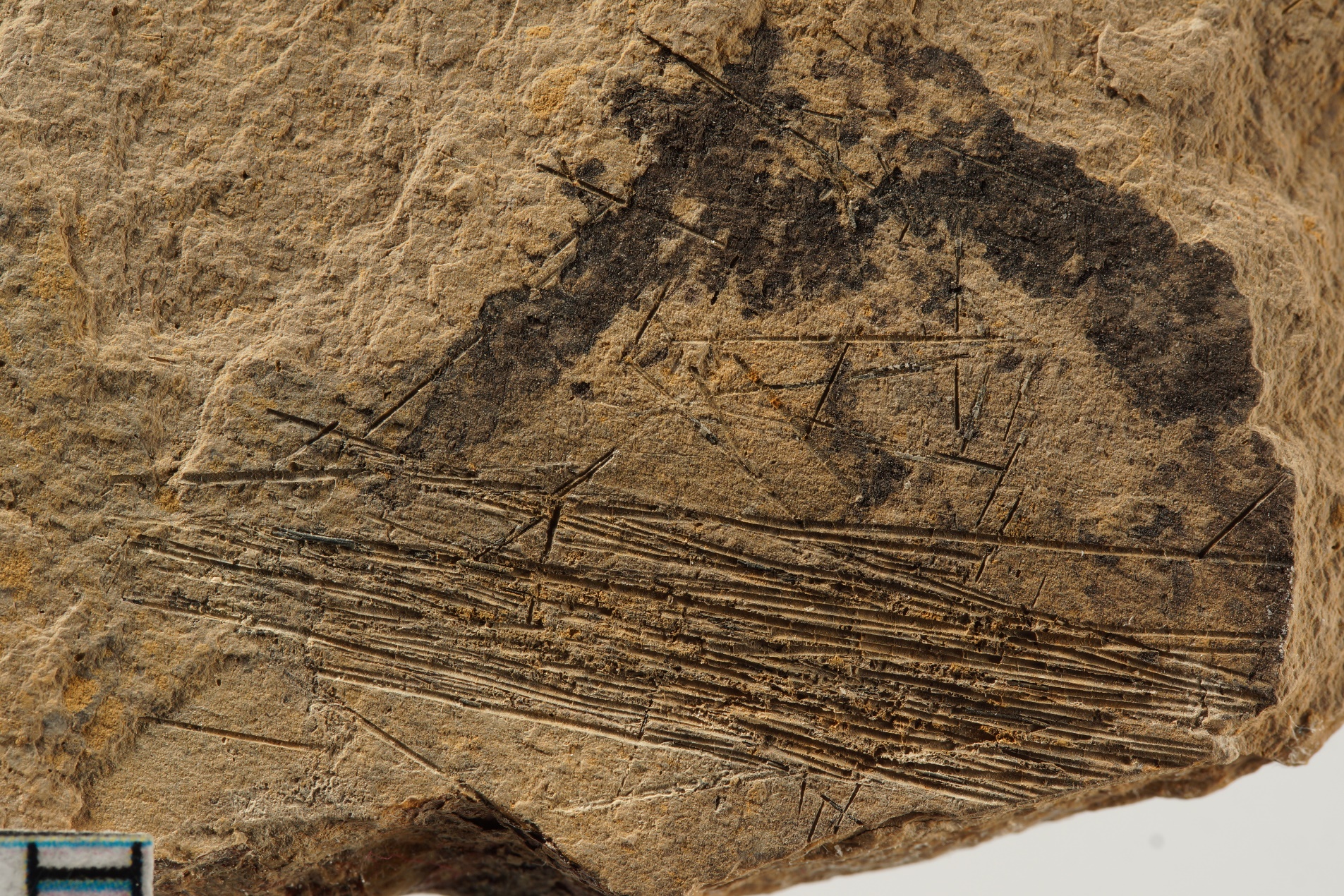

GZG.INV.1002 B

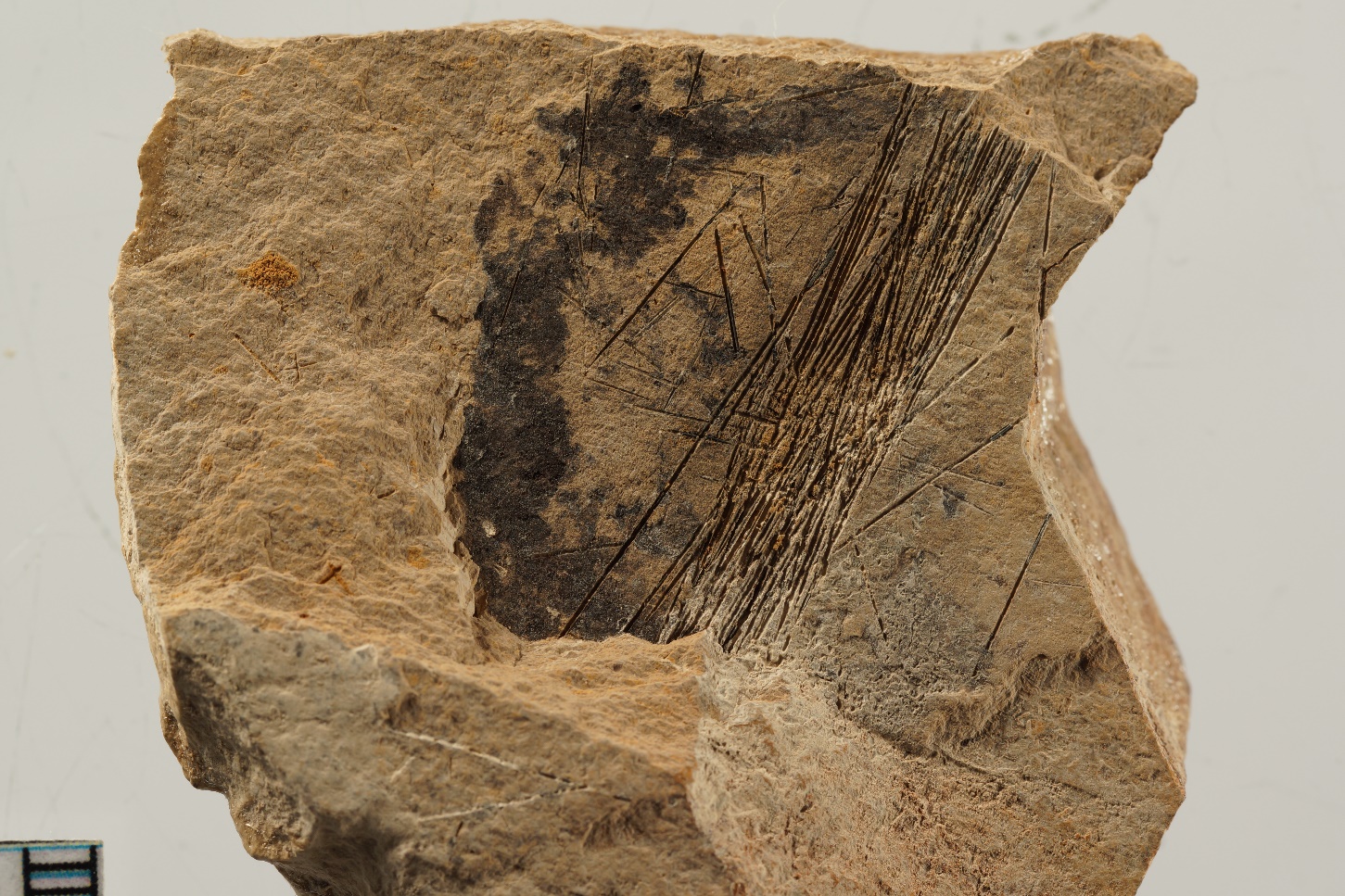

GZG.INV.998

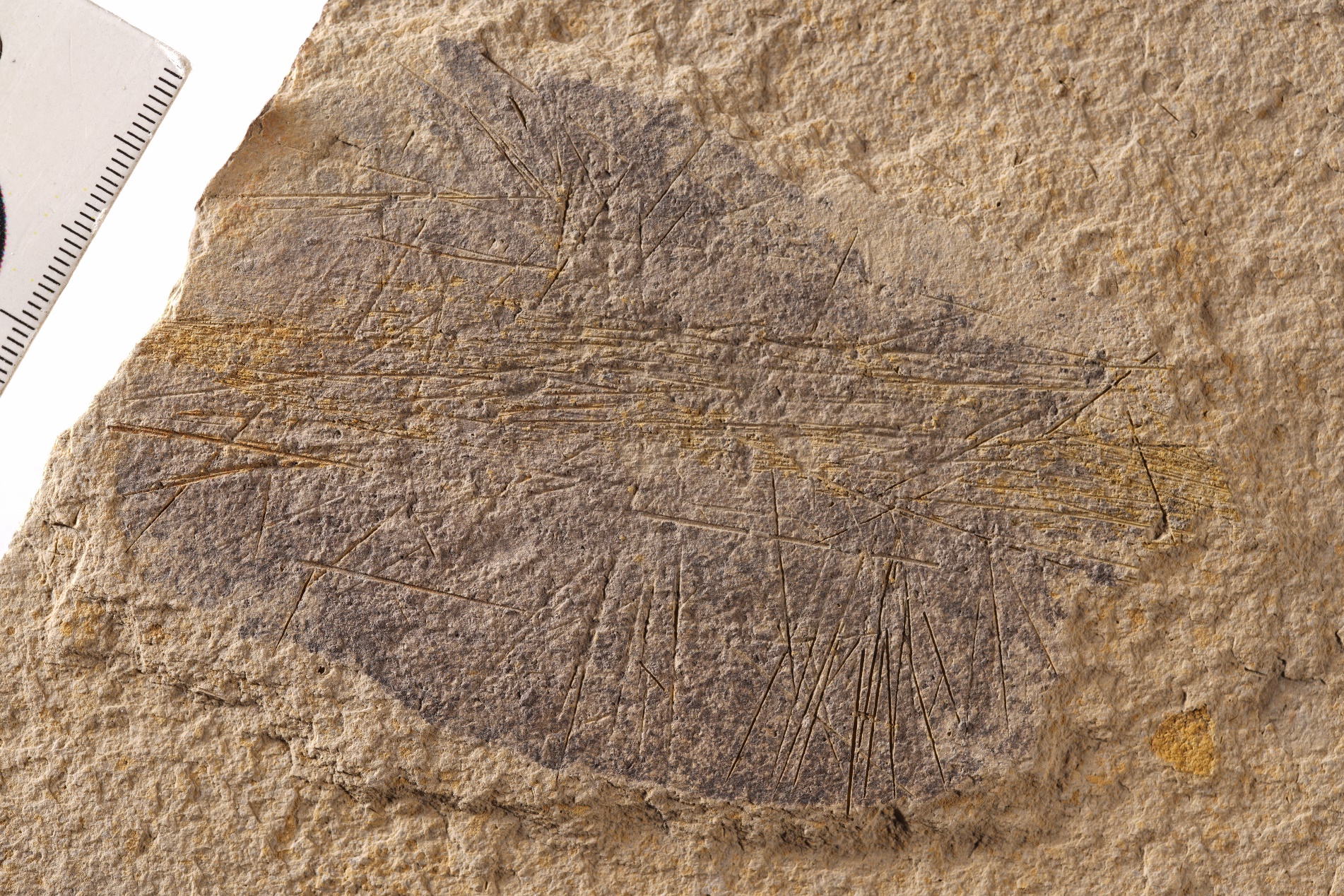

GZG.INV.1000 A

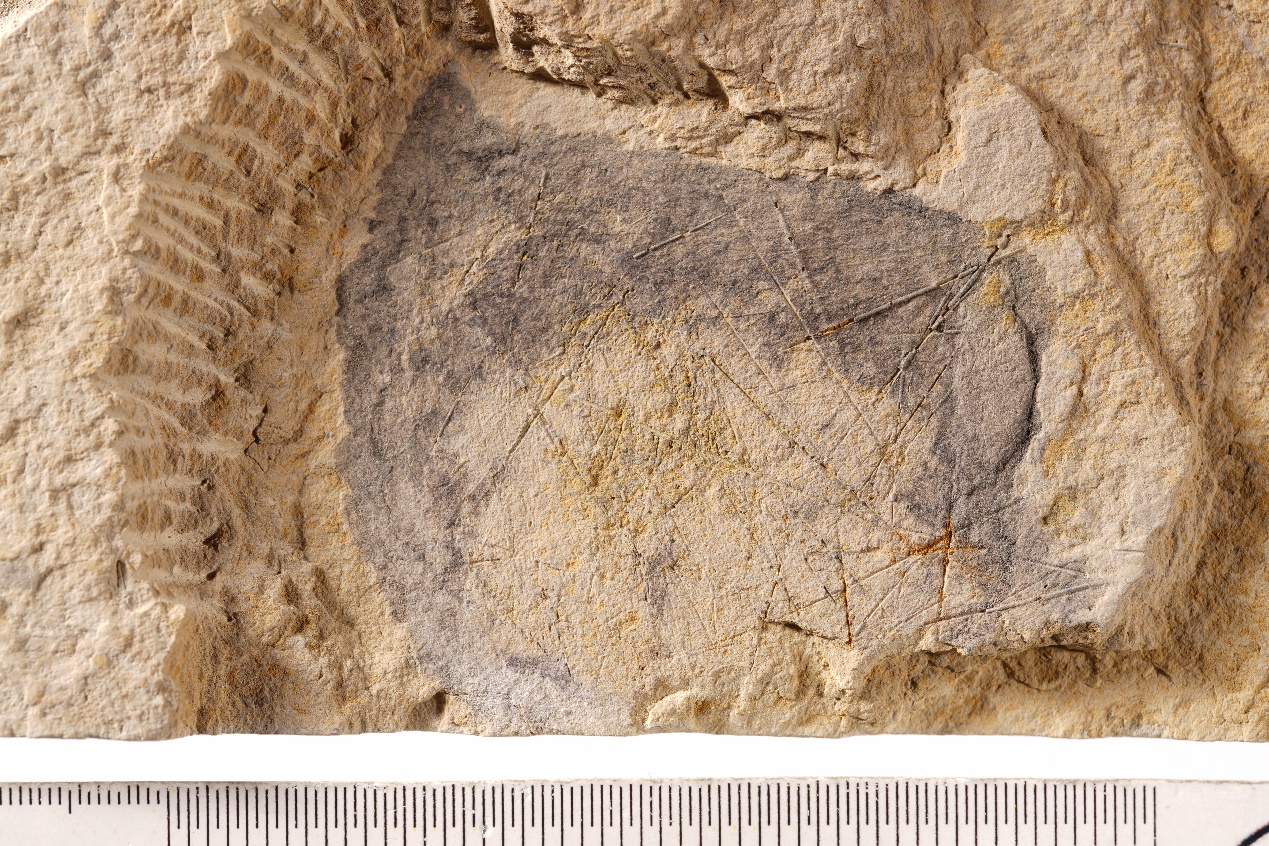

GZG.INV.1000 B

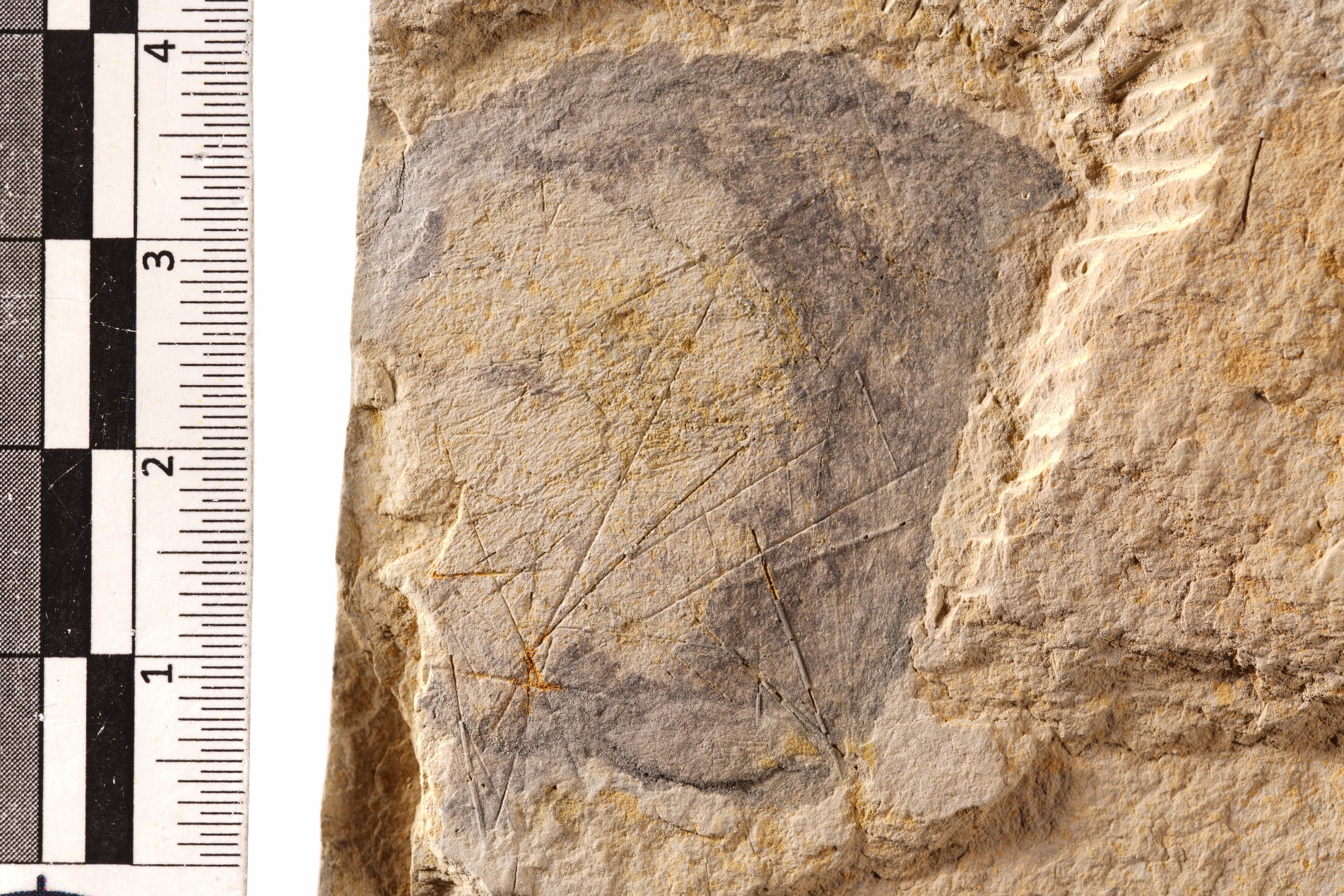

GZG.INV.1003 A

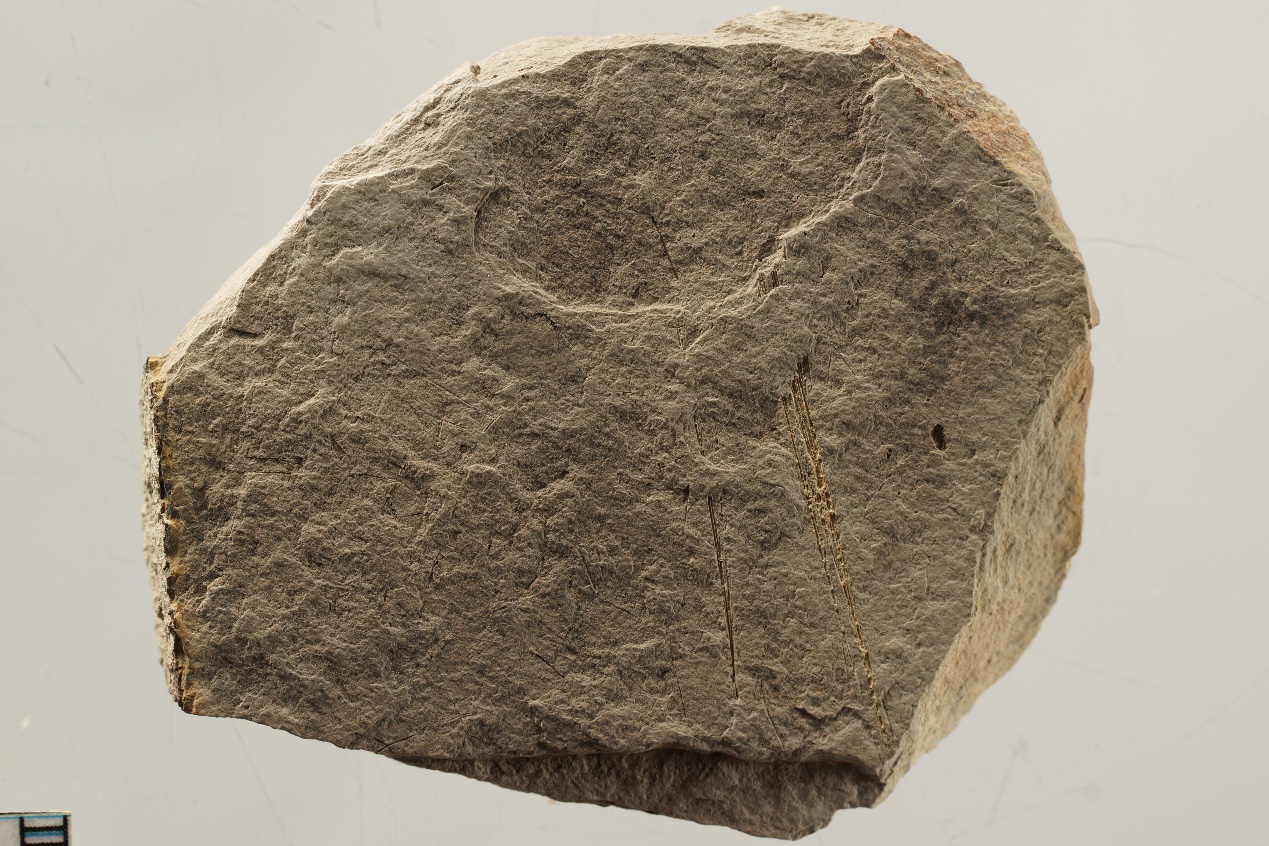

GZG.INV.1003 B

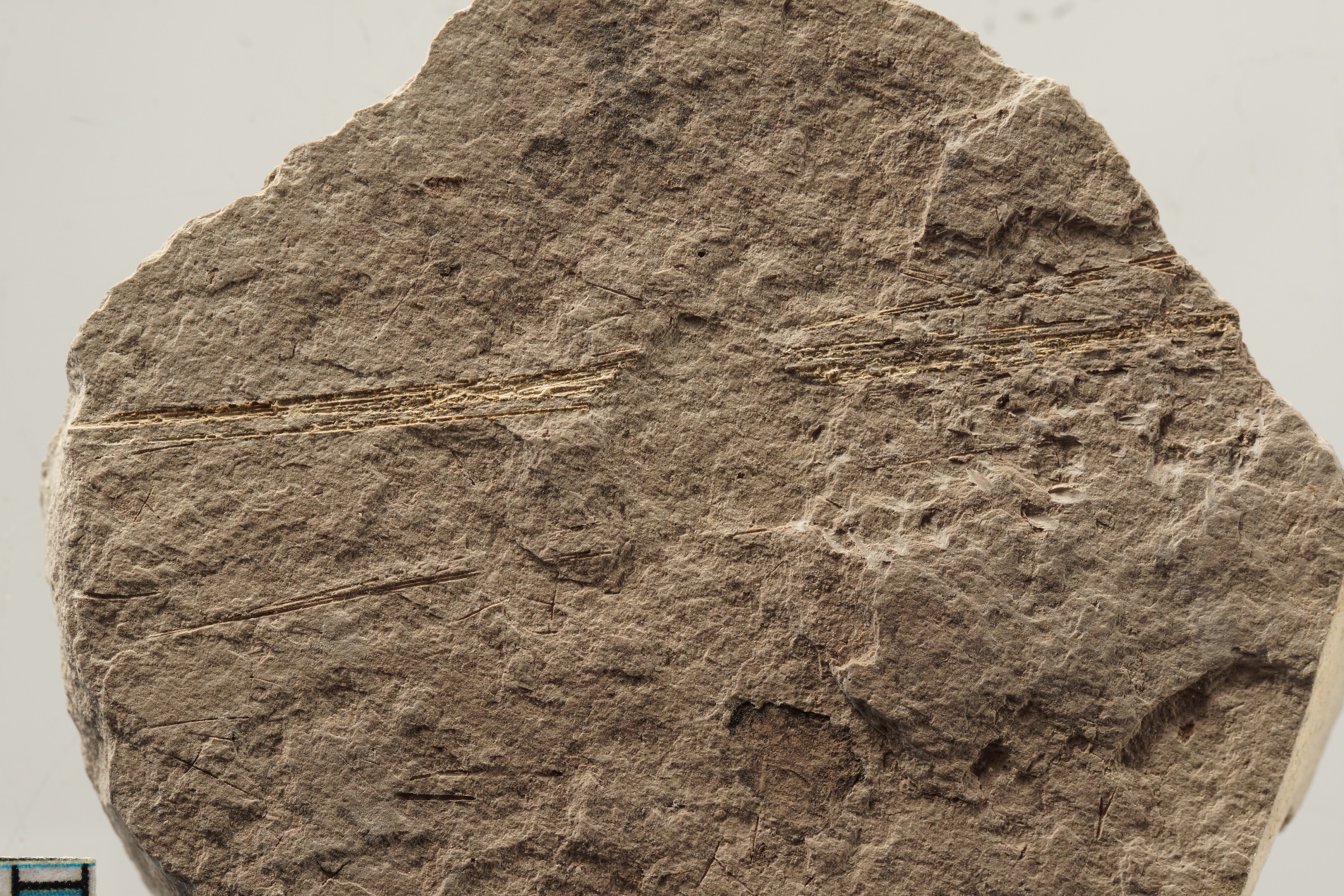

GZG.INV.1001 A

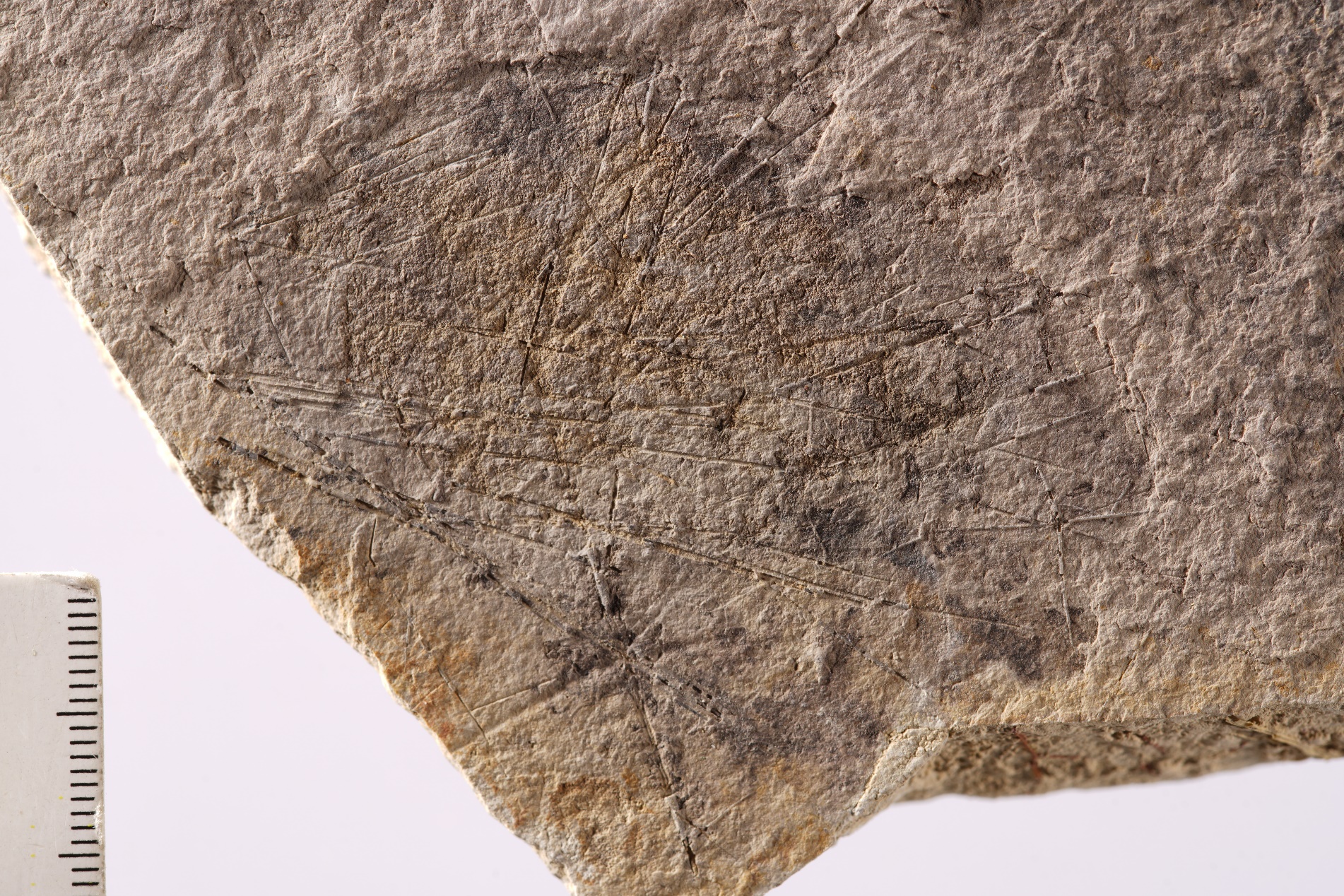

GZG.INV.1001 B

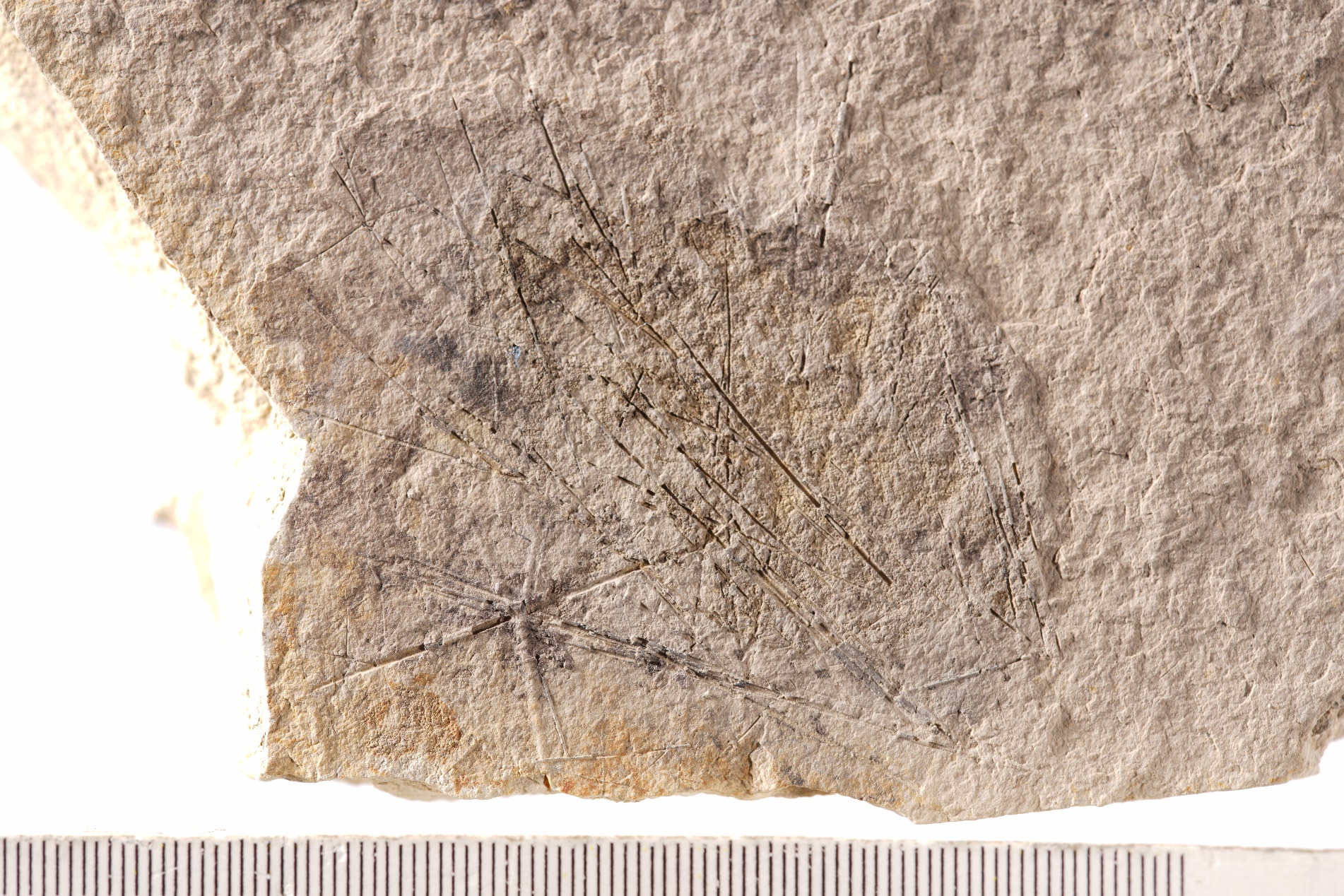

**2. X-Ray Fluorescence (XRF) results of the specimen** GZG.INV.998

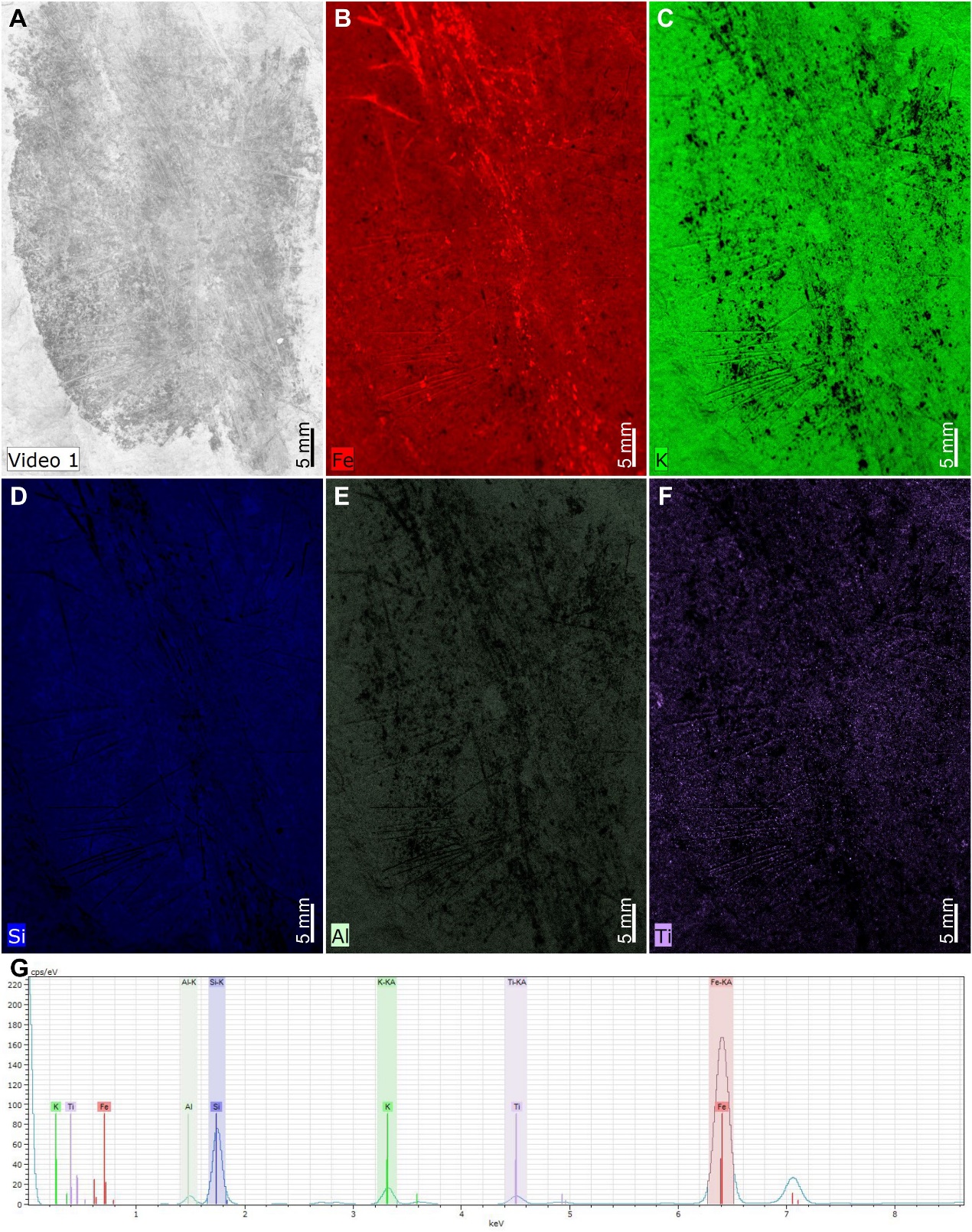

**3. Systematic palaeontology**

Phylum Porifera Grant, 1836

Class Hexactinellida Schmidt, 1870

Order and family uncertain

Genus *Hyalosinica* Mehl & Reitner in Steiner *et al*., 1993

1993 *Hyalosinica* Mehl & Reitner in Steiner *et al*., p. 305.

1993 *Solactiniella* Mehl & Reitner in Steiner *et al*., p. 309.

2005 *Hyalosinica* Mehl & Reitner; Xiao *et al*., p. 105.

2005 *Solactiniella* Mehl & Reitner; Xiao *et al*., p. 108.

2005 *Solactiniella* Mehl & Reitner; Wu *et al*., p. 1049.

2010 *Solactiniella* Mehl & Reitner; Yang *et al*., p. 353.

2016 *Solactiniella* Mehl & Reitner; Botting & Peel, p. 478.

**Type species.** *Hyalosinica archaica* Mehl & Reitner in Steiner *et al*., 1993.

**Emended diagnosis.** The sponge body is composed of an ovoid or globular main part and a long stalk-like root tuft. Spicules of the ovoid body are dominated by large diactines, which are organized as fan-shaped clusters and radiating toward and protruding beyond the margins of the body. A few small diactines, stauractines, and hexactines are scattered in the interspace of the large diactine clusters. The root tuft is interwoven by twisted bundles of long diactines (and maybe monactines). The upper end of the tuft deeply inserts into the ovoid body and the relatively loose basal end originally anchored to the seafloor.

**Remarks.** The new fossils reveal that the Sancha specimens previously classified into *Hyalosinica* and *Solactiniella* are incomplete body parts of the same species. Therefore, the genus *Solactiniella* is considered as a junior synonym of *Hyalosinica*, and the diagnosis of *Hyalosinica* is emended and expanded to describe the complete sponge body.

*Hyalosinica archaica* Mehl & Reitner in Steiner *et al*., 1993

Figures 1–3

1993 *Hyalosinica archaica* Mehl & Reitner in Steiner *et al*., p. 305, pl. 4, Fig. 1a, b.

1993 *Solactiniella* *plumata* Mehl & Reitner in Steiner *et al*., p. 309, pl. 2, Fig. 1.

?2005 *Hyalosinica* *archaica* Mehl & Reitner; Xiao *et al*., p. 107, Fig. 5D.

2005 *Solactiniella* *plumata* Mehl & Reitner; Xiao *et al*., p. 108, Fig. 9A, B.

?2005 *Solactiniella* cf. *plumata* Mehl & Reitner; Wu *et al*., p. 1049, Fig. 7.1A.

2010 *Solactiniella plumata* Mehl & Reitner; Yang *et al*., p. 353, pl. 3, Fig. 4.

?2016 *Solactiniella* cf. *plumata* Mehl & Reitner; Botting & Peel, p. 478, Fig. 8E.

?2021 Uncertain hexactinellid B; Luo *et al*., p. 81, Fig. 6B, D.

**Emended diagnosis.** As for the genus.

**Holotype.** The holotype of *Hyalosinica archaica* (San 109A, B) from the lower Niutitang Formation (basal Stage 3 of Cambrian) in Sancha section, Hunan Province, China was originally described by Mehl & Reitner in Steiner *et al*. (1993). Whereas the original holotype of *Solactiniella plumata* (San 101 A, B) described by Mehl & Reitner in Steiner *et al*. (1993) is considered as a junior synonym and hence listed as the holotype of the subgenus.

**Specimens.** GZG.INV.998 - GZG.INV.1003 from the lower Niutitang Formation (basal Stage 3 of Cambrian) in Sancha section, Hunan Province, China.

**Description.** See the main text.

GZG.INV.998 (main body + long root tuft)

GZG.INV.999 A, B (main body + long root tuft)

GZG.INV.1000 A, B (main body)

GZG.INV.1001 A, B (main body)

GZG.INV.1002 A, B (root tuft)

GZG.INV.1003 A, B (root tuft)

**4. Silhouettes of stalk-bearing sponges**

*Hyalosinica archaica* Mehl & Reitner in Steiner *et al*., 1993 (this paper)

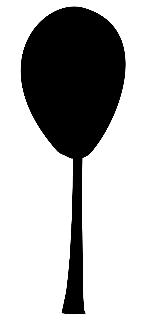

*Protospongia cyathiformis* Dawson & Hinde, 1888

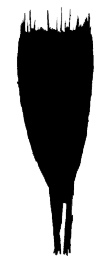

*Stioderma coscinum* Finks, 1960

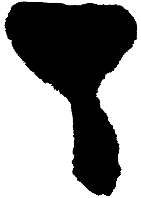

*Hyalonema (Hyalonema) sieboldii* Gray, 1835

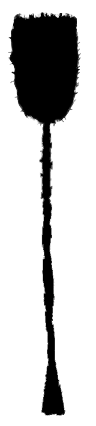
